# Mosquito-directed PROTACs to block malaria transmission

**DOI:** 10.64898/2026.04.16.719026

**Authors:** Neha Rawat, Jhalak Singhal, Bharti Goyal, Nidha Parveen, Chiging Tupe, Kanika Gupta, Soumyananda Chakraborty, Kailash Pandey, Shailja Singh

**Affiliations:** Special Centre for Molecular Medicine, Jawaharlal Nehru University, New Delhi, India; BIO-MED (P) LTD., Ghaziabad, India; ICMR-National Institute for Malaria Research, New Delhi, India; AcSIR -Academy of Scientific and Innovative Research, Ghaziabad, Uttar Pradesh, India; Birla Institute of Hyderabad Campus, Telangana, India Technology & Science, Pilani, India; School of Interdisciplinary Research, IIT-Delhi, India

**Keywords:** PROTAC, Anopheles p38 MAPK, Malaria, Vector-targeted antimalarial, Transmission blocker

## Abstract

The mosquito stage of the *Plasmodium falciparum* life cycle is an attractive target for intervention since it is crucial for the sexual reproduction and transmission of parasites to human host. Mosquito determinants crucial for parasite infection and growth pose as lucrative targets for transmission blockers. Owing to the fact that p38 MAPK has role in immune response and vector competence, we have evaluated the potential of PROTAC molecule (NR-7h) to degrade *Anopheles stephensi* p38 MAPK (*As*p38 MAPK), a conserved serine/threonine kinase involved in stress reactions, midgut homeostasis, and parasite survival. PROTAC-mediated degradation of *As*p38 MAPK led to the disrupted development of the parasite, suggesting its crucial function in vector competence. Furthermore, NR-7h-treated mosquitoes showed higher expression of immune genes such Rel-2, TEP1, APL1, and NOS, suggesting that p38 MAPK regulates host immunity in a way that promotes parasite persistence. PROTAC-mediated degradation of target proteins, provides a more persistent and resistance-proof therapeutic effect than traditional kinase inhibitors. Our findings establish PROTACs as a novel vector-targeted strategy for the development of endectocides to limit malaria transmission.

## Introduction

Malaria, caused by the protozoan parasite, *Plasmodium falciparum* (*P. falciparum*), is a worldwide public health problem, largely due to the complicated life cycle of the parasite, including two hosts, humans and *Anopheles* mosquito, and multiple life stages^[1,2]^. Until recently, control efforts for these diseases have primarily focused on killing or blocking transmission of the human host parasite e.g., antimalarial drugs^[3–5]^, vaccines^[6,7]^ rather than the mosquito vector responsible for transmitting the disease. The mosquito-stage phase is of paramount importance, because it is the only life-cycle stage where parasites sexually reproduce, recombine genetically and develop into the infective sporozoite forms.

In the case of *Anopheles* mosquitoes, the immune response against *P. falciparum* is dynamic and acts at various levels of parasite development. Following ingestion, the parasite has to overcome physical barriers like the peritrophic matrix, as also an immunological challenge of the mosquito midgut microbiota that elicits the basal immune response by the expression of antimicrobial peptides^[8]^. Humoral and cellular reactions are induced after ookinete invasion of the midgut epithelium^[9–12]^. One of the major determinants of humoral immunity is the TEP1, a complement-like protein that along with the co-factors LRIM1 and APL1C, recognizes and kills parasites^[13,14]^. Hemocytes, the major cellular immune component, participate in cellular defense through phagocytosis, melanization and release of ROS^[15,16]^. Many evolutionarily conserved signaling cascades trim these responses. Early activation of the Toll and Imd signaling cascades regulates expression of antimicrobial peptides (AMPs) and effector molecules^[17–19]^. The JNK and STAT pathways act later on, specifically during oocyst development, by modulating nitric oxide synthase (NOS) and other immune genes^[20,21]^. Earlier evidence additionally underlines the role of the p38 Mitogen-activated protein kinase (MAPK) pathway, such as the *As*p38 MAPK isoform, in favouring mosquito survival after parasite challenge. This survival advantage likely reflects the greater co-evolutionary history between mosquitoes and *Plasmodium* than with vertebrate hosts, allowing mosquitoes to withstand and accommodate chronic infections^[22]^. These immune mechanisms not only restrict parasite growth but also can identify potential determinants for vector-targeted transmission-blocking interventions to control malaria.

The p38 mitogen-activated protein kinase (MAPK) pathway is a conserved signal cascade that is involved in cellular responses to stress, inflammation, and pathogen infection in various organisms^[23]^. p38 MAPK plays a vital role in mediating innate immune reactions and physiological homeostasis in disease vectors like mosquitoes^[24]^, ticks^[25]^, and sandflies^[26]^ during pathogen challenge. In *Anopheles* mosquitoes, p38 MAPK signaling is triggered by infection with *P. falciparum* and has been reported to facilitate vector survival through the regulation of oxidative stress and anti-apoptotic pathways, possibly enabling successful parasite development and transmission^[24]^. In the vector *Ixodes scapularis*, the Lyme disease vector, p38 signaling regulates tick immune response and persistence of the pathogen^[27,28]^. Additionally, the route is linked to hemocyte function, gut barrier integrity, and vector responses to microbial disturbance^[29]^. These results show how p38 MAPK has developed into an essential modulator of interactions between vectors and pathogens. By impairing vector competence without directly harming the pathogen, targeting this pathway is a viable strategy to stop the spread of the infection and lower the likelihood that resistance will emerge.

Proteolysis-targeting chimeras (PROTACs) are a novel class of heterobifunctional chemicals designed to target protein breakdown by means of the ubiquitin–proteasome system (UPS). A PROTAC molecule is composed of two covalently bonded moieties: one for binding a target protein and another for drawing in an E3 ubiquitin ligase, which enables ubiquitination and proteasomal degradation of the target protein^[30,31]^. PROTACs catalytically and selectively eliminate disease-associated proteins, including those that are “undruggable” due to their poor ligandability or lack of enzymatic activity, in contrast to traditional small-molecule inhibitors, which frequently work by temporarily inhibiting enzymatic activity^[21,32]^. This approach has shown promise as a treatment for a variety of clinical conditions, such as inflammatory diseases, cancer, and neurodegeneration^[33]^. Interestingly, application of PROTACs to target proteins in disease vectors remains underexplored. In malaria, a PROTAC-based method for the selective protein degradation of mosquito p38 MAPK may impede the growth of parasites and disrupt their transmission. By focusing on vector-stage biology, this approach presents a promising way to supplement current antimalarial efforts and could be a novel intervention in transmission-blocking control of malaria.

## Methods

### Sequence analysis and phylogeny

To identify homologs of human p38 MAPK and Cereblon in *Anopheles stephensi*, the amino acid sequences of *Homo sapiens* p38 MAPK (HsMAPK14; UniProt ID: Q16539) and Cereblon (HsCRBN; UniProt ID: Q96SW2) were used as query sequences in BLAST searches against the *Anopheles stephensi* genome database available at VectorBase (https://vectorbase.org/vectorbase/app/). Similarly, homologs of DDB1 and RBX1 in *A. stephensi* were identified using the *H. sapiens* DDB1 (UniProt ID: Q16531) and RBX1 (UniProt ID: P62877) amino acid sequences, respectively. For the identification of *A. stephensi* cullin homologs, amino acid sequences of *H. sapiens* cullins—CUL1 (Q13616), CUL2 (Q13617), CUL3 (Q13618), CUL4A (Q13619), CUL4B (Q13620), CUL5 (Q93034), CUL7 (Q14999), and CUL9 (Q8IWT3)— were used as queries in BLAST searches against the *A. stephensi* genome database in VectorBase. Sequence alignment was performed using Clustal Omega (https://www.ebi.ac.uk/Tools/msa/clustalo/)^[34]^, and amino acid sequences were analysed for conserved domains and motifs using CD-BLAST program (https://www.ncbi.nlm.nih.gov/Structure/cdd/wrpsb.cgi). 3-D structures of both the proteins were obtained from the AlphaFold Database and aligned using PyMol software. For phylogeny, amino acid sequences were aligned using the Molecular Evolutionary Genetic Analysis (MEGA12) program (https://www.megasoftware.net/), the phylogenetic tree was also constructed using MEGA12 software^[35]^. The components of *A. stephensi* E3 ligase complex were aligned over the CRL4-CRBN complex of humans by PyMol software.

### Molecular Docking

The *in silico* molecular docking analysis between *As*p38 MAPK and PH-797804 inhibitor was done using CB-DOCK2 server (https://cadd.labshare.cn/cb-dock2/index.php)^[36]^.

### Immunofluorescence Assay

Midguts from dissected mosquitoes were washed, fixed in 4% paraformaldehyde and permeabilized in 0.01 % Triton X-100. Subsequently, the midguts were stained with p38 MAPK antibody (1:100; Biolegend, #BF8015), followed by staining with anti-mouse Alexa 488 antibody (1:100; Invitrogen, #A11029) for 1 hour. After washing, the midguts were mounted with Anti-fade solution supplemented with DAPI (Sigma, #F6057) and visualized under fluorescence microscope at 10X magnification.

### Mosquito rearing, drug treatment and *Plasmodium >berghei* infection

For the experiments, *A. stephensi* were reared and maintained at 28°C ± 2, RH = 80% in the insectary fitted with a simulated dawn and dusk machine with a 12 h Light/Dark cyclic period at ICMR-NIMR. All protocols for rearing and maintenance of the mosquito culture were approved by the ethical committee of the ICMR-NIMR. For drug treatment, 4–5-day old laboratory reared *A. stephensi* mosquitoes were starved and kept on water overnight and then allowed to feed on NR-7h PROTAC through membrane feeding system for 30 minutes. After 10 hours, the mosquitoes were then allowed to feed on *Plasmodium berghei (P. berghei)*-infected mice with 4-5% gametocytes. Post 9-10 days of *P. berghei* infection, the mosquitoes were dissected and midguts were collected. Midguts were then washed and stained with 0.2% mercurochrome (Sigma-Aldrich) to determine the oocyst count. The stained midguts were visualised at 10X magnification under compound microscope.

### Quantitative real-time PCR (qRT-PCR)

4-5 days old female *Anopheles* mosquitoes were treated with 10μM NR-7h and allowed to feed on *P. berghei* infected mice. Mosquitoes were collected at two time points - 24 hours post infection (24 h.p.i) and 8 days post infection (8 d.p.i) for analysis of immune gene expression while for determination of parasitic load the midguts were collected post 8 days of *P. berghei* infection. For the determination of p38 MAPK transcript expression in untreated and NR-7h treated mosquito midguts, the midguts were dissected at 24 h.p.i. For RNA isolation, 25 midguts were dissected and homogenized in Trizol reagent (Sigma-Aldrich; T9424). cDNA was synthesized from 1000ng of isolated RNA by iScript™ cDNA Synthesis Kit (Bio-Rad; #1708891). Expression levels of *As*p38 MAPK, *As*Cullin 4B, *As*RING-box protein 1 (RBX1), *As*DNA-damage binding protein (DDB1), *As*Cereblon, TEP1, Rel-2, Defensin, Gambicin, Cecropin C, LRIM1, APL1, NOS, LRRD7/LRIM17 and *Pb*18S rRNA were analysed using PowerUp SYBR Green PCR Master Mix (Thermo Fisher Scientific #A25742) on Applied Biosystems Real-Time PCR System (ABI). The expression levels of selected genes were normalized to Ribosomal protein S7 (RPS7) and calculated by 2^−ΔΔCt^ method.

The primers sequences are: *Asp38 MAPK* (FP: 5’-ATCATTCACCGGGATTTGA-3’;RP:5 ‘-GTGCATCCAGTTGAGCAT-3’), *AsCullin4B* (FP: 5’-AGTTAATTAACGAACACGTAAC-3’; RP: 5’GTTACGTGTTCGTTAATTAACT -3’), *AsRBX1*(FP: 5’-ATGGATATCGACGAGGAAGAGTTC-3’; RP: 5 ‘-CTAATGGCCGTACTTCTGGAA -3’), *AsDDB1*(FP: 5’-ATGAAATCAGCCTGAAGGAT-3’; RP: 5 ‘-GCGTAGCAGTTAATCGTGC -3’), *AsCereblon*(FP: 5’-AGTGTATCCGGGCGAAATAG-3’; RP: 5 ‘-CCGAACGATGTAGAACCGT - 3’), *TEP1*(FP: 5’-CGAGAAGCATCTGATAAATAAAGC-3’; RP: 5 ‘-TACCAGAGCGGTTACGAAGAT - 3’), *Rel-2*(FP: 5’-AGACTACGGTAAACCGCATC-3’; RP: 5 ‘-CGGACTTGATCACAAGATGAT - 3’), *Defensin* (FP: 5’-GTCGTGGTCCTGGCGGCTCT-3’; RP: 5 ‘-ACGAGCGATGCAATGCGCGGCA - 3’), *Gambicin* (FP: 5’-ATGAAGCAAGTGTGCATTGTTCT-3’; RP: 5 ‘-TCACAAGAAGCACTCCGTAATG - 3’), *Cecropin C* (FP: 5’-ATGAACTTCAAGCTGCTCTTTC-3’; RP: 5 ‘-CTATCCAAGGACACCCTTTACAC - 3’), *LRIM1*(FP: 5’-GCGTCGGTTCGGAAAAGGAGCGG-3’; RP: 5 ‘-TACATATCCCAATCGCGGATGGC - 3’), *APL1*(FP: 5’-AGAGTCGGCAGGCGTTCAA-3’; RP: 5 ‘-GCTTGTCGGTCTTCAGGGTCAG - 3’), *NOS*(FP: 5’- GACCAAACCGGTCATCCTGAT-3’; RP: 5 ‘-GGAATCTTGCAGTCAACCATTTC - 3’), *LRRD7/LRIM17*(FP: 5’-AAGCTGATCACACTCGATCTGT-3’; RP: 5 ‘-TACGCACCATCACCGGGAACGA - 3’), *RPS7* (FP: 5’-GAAGGCCTTCCAGAAGGTACAGA-3’; RP:5’-CATCGGTTTGGGCAGAATG-3’) and *Pb18S rRNA* (FP:5’-GGAGATTGGTTTTGACGTTTATGTG-3’; RP: 5 ‘- AAGCATTAAATAAAGCGAATACATCCTTAC - 3’)

### Data analysis

Data are presented as mean ± standard deviation (SD). Each experiment was independently repeated three times using different batches. Graphs and corresponding statistical analyses were generated using GraphPad Prism (GraphPad Software, USA). Statistical significance was assessed using an unpaired t-test with Welch’s correction. A p-value of less than 0.05 was considered statistically significant. Significance levels are indicated as follows: *p < 0.05, **p < 0.01, ***p < 0.001, and ****p < 0.0001.

## Results

### *Anopheles stephensi* genome encodes for components of CRL4 E3 ligase complex

Among the multiple E3 ligases in eukaryotes, Cullin-RING Ligases (CRLs) represent the largest family of E3 ubiquitin ligases, built around a Cullin scaffold protein. They assemble into muti-subunit complexes where the Cullin acts as a structural backbone, the RING protein (RBX) recruits and activates the ubiquitin-loaded E2 enzyme, adaptor protein and a substrate adaptor molecule that recognizes specific target proteins. The Cullin isoforms vary between the eukaryotic organisms, for instance, *Homo sapiens* genome encodes for 8 Cullin isoforms namely Cullin-1,2,3,4A,4B,5,6,7,and 9^[37]^, whereas *Plasmodium falciparum* (*P. falciparum*) has only two Cullin isoforms – *Pf*Cullin-1 and *Pf*Cullin-2^[38]^.*A. stephensi* expresses five cullin isoforms namely Cullin-1 (*As*Cullin1, ASTEI20_038001), Cullin-2 (*As*Cullin, ASTEI20_037441), Cullin-3 (*As*Cullin, ASTEI20_033142), Cullin-4B (*As*Cullin4B, ASTEI20_034952) and Cullin-5 (*As*Cullin5, ASTEI20_031831). *As*Cullin4B shows 65% sequence similarity to Human Culllin-4A (*Hs*Cullin4A) and Human Cullin-4B and has the typical domain architecture of cullin proteins including the cullin homology domain and the neddylation motif (**Figure 1a)**. Cullin4A and Cullin4B associates with RBX1, DNA damage binding protein (DDB1) and DDB1- and CUL4-associated factor (DCAF) to form a functional Cullin4-E3 ligase complex. *A*.*stephensi* also encodes for RBX1A (*As*RBX1A, ASTEI20_038593) which has 84% sequence identity with Human RBX1 (*Hs*RBX1) with the signature C3H2C2 zinc binding domain^[39]^**(Figure 1b)**. The DDB1 protein of *A*.*stephensi* (AsDDB1, ASTEI20_032418) has 67% sequence identity to Human DDB1 (*Hs*DDB1) **(Figure 1c)**. Lastly, the *A. stephensi* encodes for multiple DCAFs and a DCAF-type adaptor molecule, Cereblon (*As*Cereblon, ASTEI20_034852) with 37% sequence similarity to Human Cereblon (*Hs*Cereblon) and a thalidomide binding C-terminal domain **(Figure 1d)**. The substrate adaptor molecule in the E3 ligase complex is specific to each Cullin isoform e.g., Skp-1-F-box in CRL1/SCF^[40]^, Elongin B/C-SOCS/VHL in CRL2^[41]^, and DDB1-DCAF in CRL4^[42]^. Additionally, expression of Cullin4B, RBX1, DDB1, and Cereblon transcripts was assessed by RT-PCR using whole *Anopheles* mosquito- and midgut-derived cDNA as template. Gene-specific primers were designed, and amplification confirmed the presence of corresponding transcripts (**Figure 1e.)**. Furthermore, all the components were structurally aligned and superimposed onto the resolved human CRL4 E3 ligase complex **(Figure 1f)**^[43]^ to generate a structural model of the Anopheles CRL4-CRBN E3 ligase complex. **(Figure 1g)**. The expression of components suggests *that A*.*stephensi* has a CRL4-CRBN complex similar to *H. sapiens* which can be exploited for targeted degradation of proteins by PROTACs.

**Table 1:** Comparative Sequence Identity of CRL4 E3 Ligase Components Between Human and *Anopheles stephensi*.

| CRL Component | Human Protein | *A. stephensi* Protein | % Sequence Identity |
| --- | --- | --- | --- |
| **Cullin** | CUL4A | Cullin1 | 33 |
|  | CUL4A | Cullin2 | 28.43 |
|  | CUL4A | Cullin3 | 37.62 |
|  | CUL4A | Cullin4B | 65.41 |
|  | CUL4A | Cullin5 | 25.44 |
|  | CUL4B | Cullin1 | 33.14 |
|  | CUL4B | Cullin2 | 29.15 |
|  | CUL4B | Cullin3 | 38.78 |
|  | CUL4B | Cullin4B | 65.07 |
|  | CUL4B | Cullin5 | 26.16 |
| **RING Protein** | RBX1 | RBX1 | 84 |
| **Adaptor** | DDB1 | DDB1 | 67 |
| **Substrate Receptor** | CRBN | CRBN | 37 |

**Figure 1.**
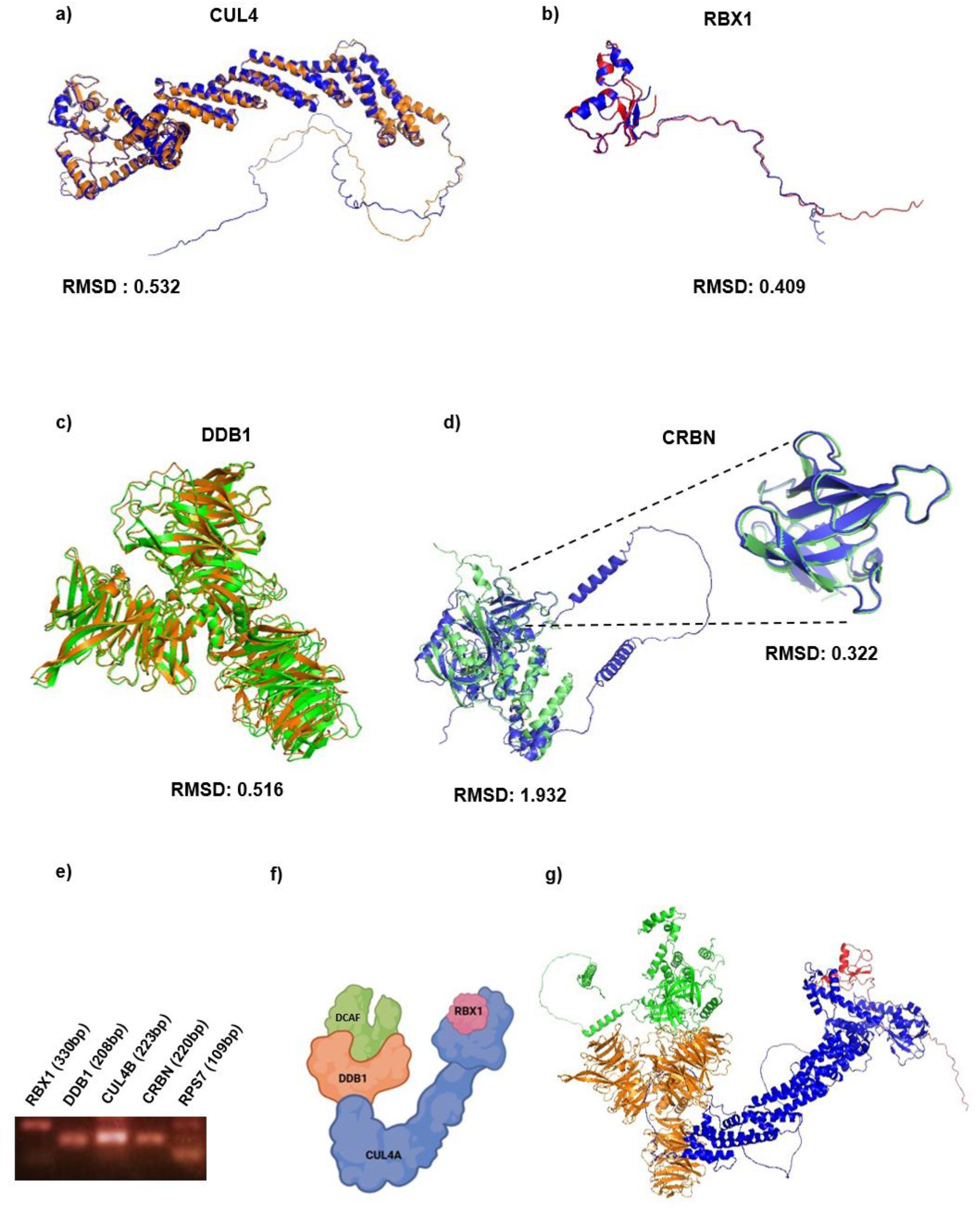
Components of CRL4 E3 ligase complex in *Anopheles stephensi*. Structural alignment of *Anopheles stephensi* CRL4 E3 ligase components with Human homologues **(a)** Cullin 4B (CUL4B; *As*CUL4B-blue, *Hs*CUL4A-orange), **(b)** Ring Box Protein 1 (RBX1; *As*RBX1-red, *Hs*RBX1-blue), **(c)** Cereblon (CRBN; *As*CRBN-blue, *Hs*CRBN-green) & the thalidomide binding domain and **(d)** DNA Damage binding Protein 1(DDB1; *As*DDB1-orange, *Hs*DDB1-green) with respective RMSD values. **(e)** CRL4-DCAFE3 complex model in humans, and **(f)** Modelled structure of CRL4B-CRBN Complex of identified components of *A*.*stephensi*

### p38 MAPK is conserved among *Anopheles* species

The domain analysis of *As*p38 MAPK (ASTEI06041) revealed the presence of conserved kinase motifs including VAIK, HRD, DGF and TXY motifs **(Figure 2a)**. VAIK motif is essential for ATP binding where the lysine (K) in the motif interacts with phosphates of ATP and helps in nucleotide positioning^[40]^. HRD motif is present in the catalytic site and is involved in phosphoryl transfer and substrate recognition^[41]^. DFG motif, present in activation loop, is involved in binding the divalent cations (Mg^2+^ or Mn^2+^) that are required for coordinating the phosphates of ATP necessary for catalysis^[42]^.TXY motif is located within the activation segment of MAPK and the phosphorylation of both the threonine and tyrosine residues by upstream MAP kinase kinases (MKKs) induces a conformation shift that stabilizes the active site, aligns the catalytic residues, and permits efficient substrate access^[43,44]^. The predicted protein comprises 366 amino acids and contains functional domains such as the ATP-binding site, activation loop, kinase-interacting motif (KIM) docking site, active site, and polypeptide substrate binding site, indicating strong conservation of kinase structural features across species. Multiple sequence alignment of p38 MAPK proteins from various *Anopheles* species showed highly conserved residues, particularly within the activation loop (HRD and TXY motifs) and substrate recognition regions **(Figure 2b)**. This suggests that the core catalytic and regulatory functions of p38 MAPK are maintained across mosquito vectors, underlining its functional importance in signal transduction. Phylogenetic reconstruction demonstrated a close evolutionary relationship of *As*p38 MAPK with other *Anopheles* species, clustering distinctly from outgroup species **(Figure 2c)**. Broader phylogenetic comparisons positioned mosquito p38 MAPKs within a clade separate from vertebrate MAPKs, *Plasmodium falciparum*, and other dipteran insects such as *Drosophila melanogaster* **(Figure 2d)**, indicating lineage-specific diversification while retaining conserved kinase features. Structural superimposition of *A. stephensi* and human p38 MAPK revealed a high degree of similarity, with an RMSD of 0.642 **(Figure 2e)**. This suggests strong structural conservation of the kinase, consistent with the high sequence conservation observed in catalytic motifs. Molecular docking analysis between PH-797804 (p38MAPK inhibitor), used in p38MAPK PROTAC (NR-7h), and *As*p38 MAPK revealed that the PH-797804 inhibitor binds to Val26, Val34, Ala47, Ile48, Lys49, Lue71, Ile80, Leu82, Leu101, Val102, Thr103, His104, Leu105, Met106, Gly107, Ala108, Asp109. Ala154 and Leu164 residues of *As*p38 MAPK with a binding energy of -8 kcal/mol. These residues lie within the active site of *As*p38 MAPK indicating the inhibitor targets the active site of the *As*p38 MAPK **(Figure 2f)**. Due to high similarity between p38 MAPKs of *H. sapiens* and *A. stephensi*, the PROTACs against human p38 MAPK and antibodies against human p38 MAPKs were used in the study.

**Figure 2.**
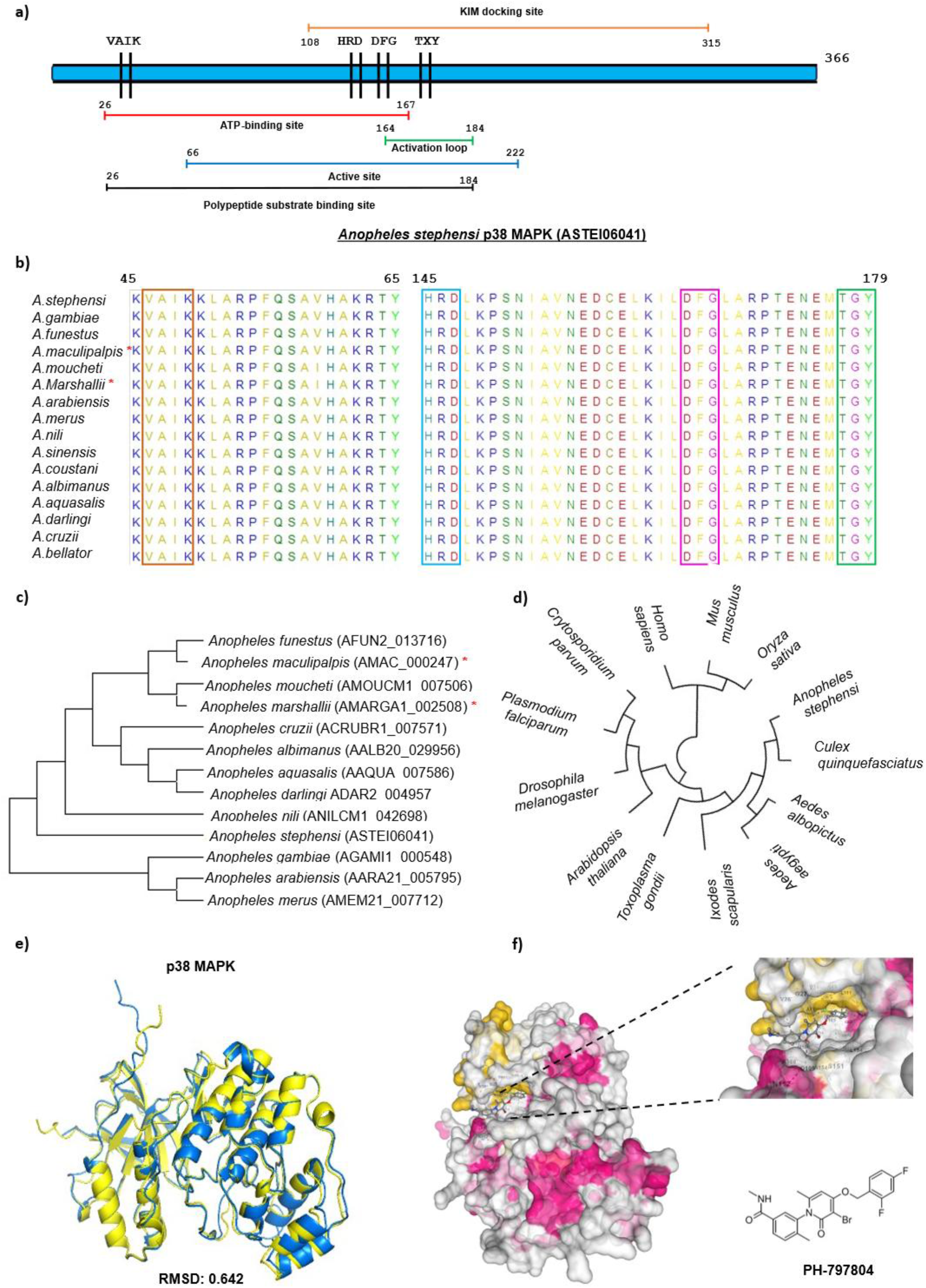
Expression of *Anopheles stephensi* midgut p38 MAPK. **(a)** Domain architecture of Anopheles stephensi p38-MAPK. **(b)** Multiple sequence alignment of p38-MAPK of Anopheles species with highlighted VAIK, HRD, DFG and TXY motifs. **(c)** Phylogenetic analysis of p38 MAPK in *Anopheles* species. **(d)** Phylogenetic tree of p38-MAPK showing evolutionary relationships among selected eukaryotic species. **(e)** Structural alignment of p38 MAPK of *Anopheles stephensi* and *Homo sapiens* with RMSD value. **(f)** Docked structure of *As*p38 MAPK and PH-797804 (p38MAPK inhibitor). (* - *Anopheles* spp. not implicated in malaria transmission)

### NR-7h PROTAC degrades *Plasmodium* infection-induced p38 MAPK in *Anopheles* midgut

To determine the expression of *As*p38 MAPK in mosquito midguts during *Plasmodium* infection, dissected midguts from uninfected and *P. berghei*infected mosquitoes were stained with human p38 MAPK-specific antibody. Immunofluorescence assay (IFA) revealed ∼2-fold increase in *As*p38 MAPK levels of *P. berghei*-infected midguts compared to uninfected midguts **(Figure 3ai)**. Mean fluorescence intensity of p38 MAPK in *Anopheles* midguts of uninfected and *P. berghei*-infected group has been shown **(Figure 3aii)**. Further, the efficacy of NR-7h PROTAC to degrade *As*p38 MAPK in midguts was also analysed in NR-7h treated dissected midgut. Immunofluorescence assay showed significant reduction (∼2-fold) in the levels of p38-MAPK **(Figure 3bi)** in NR-7h treated dissected midguts compared to untreated control. Mean fluorescence intensity of p38 MAPK in *Anopheles* midguts of untreated and NR-7h treated group has been shown **(Figure 3bii)**. Additionally, the expression of p38 MAPK transcript was also analysed in untreated and NR-7h treated midguts by qRT-PCR using midgut-derived cDNA as template **(Figure 3biii)**. The NR-7h PROTAC-treated mosquitoes were subsequently infected with *P. berghei* and post 8 days of *P. berghei-* infection (8 d.p.i), the expression of *As*p38 MAPK was also analysed in *P. berghei-*infected mosquito midguts. IFA analysis showed that the expression of *As*p38 MAPK was significantly lower (∼2-fold) in NR-7h treated midguts in comparison to untreated midguts **(Figure 3ci)**. Mean fluorescence intensity of p38 MAPK in *Anopheles* midguts of untreated and NR-7h treated group has been shown **(Figure 3cii)**.

**Figure 3.**
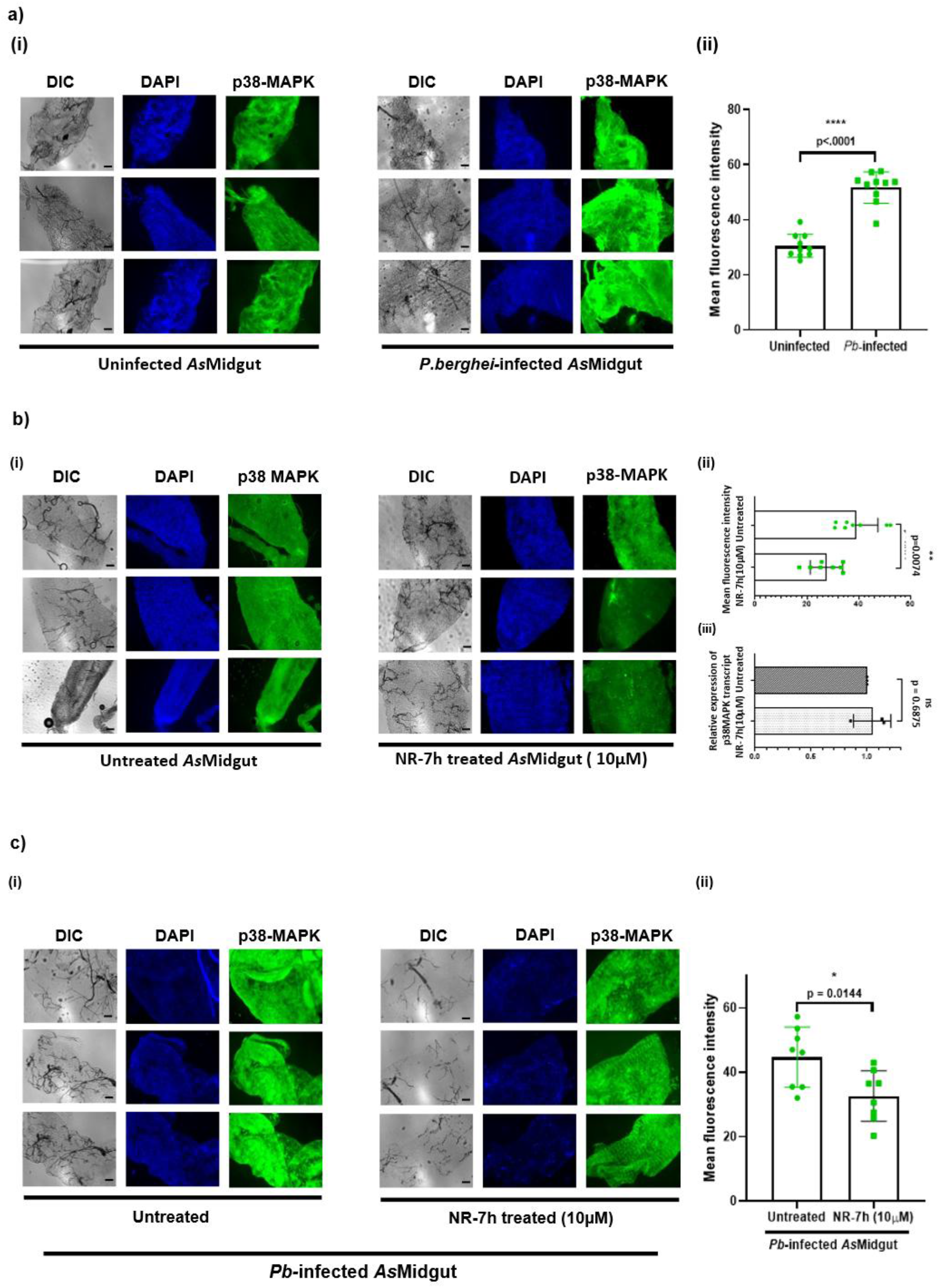
Expression of p38 MAPK in Anopheles midgut during *Plasmodium* infection and degradation of *As*p38 MAPK by NR-7h PROTAC. **(a)** (i) p38 MAPK expression in uninfected and *Plasmodium berghei (Pb)* infected dissected mosquito midgut. (ii) Mean fluorescence intensity of p38 MAPK in uninfected and *Pb*-infected midguts. **(b)** (i) p38 MAPK expression in untreated and NR-7h treated (10 µM) dissected mosquito midgut. (ii) Mean fluorescence intensity of p38 MAPK in untreated and NR-7h treated midguts (iii) Expression of p38 MAPK transcript in untreated and NR-7h treated midguts. **(c)** (i) p38 MAPK expression in untreated and NR-7h treated (10 µM) dissected *Pb*-infected mosquito midgut (8dpi). (ii) Mean fluorescence intensity of p38 MAPK in untreated and NR-7h treated *P. berghei-*infected midguts (8d.p.i).(Scale bar: 100μm)

### Targeted degradation of p38-MAPK in *Anopheles stephensi* midgut inhibits parasite infection and growth

Post treatment with 10 µM NR-7h PROTAC for 10 hours, mosquitoes were infected with *P. berghei (Pb)* by direct feeding on mice. Subsequently, parasitic load and development was determined in mosquitoes of both control and NR-7h treated group. Mosquitoes treated with NR-7h showed reduction in both infection intensity and oocyst load per midgut, where presence of at least one oocyst in mosquito midgut was considered as infection and the total number of oocysts per midgut was calculated for mean oocyst count. An infection rate of 85.4% was observed in control untreated group whereas the NR-7h treated group had an infection rate of 38%, corresponding to ∼2-fold reduction in the infection intensity **(Figure 4b)**. Similarly, significant reduction (∼ 8-fold) in oocyst load per midgut was observed in NR-7h treated mosquitoes with the value of 6.9 compared to the mean oocyst count of 56.42 in control mosquitoes **(Figure 4c and e)**. To validate the findings, the parasitic load was also determined by measuring the relative fold change in the expression of *P. berghei* 18S rRNA (*Pb*18SrRNA) in the midguts of mosquitoes of each group by qRT-PCR **(Figure 4d)**. A reduction of ∼6 fold in the expression of *Pb*18SrRNA was observed in midguts of NR-7h treated mosquitoes, which correlated with the oocyst counting data.

**Figure 4:**
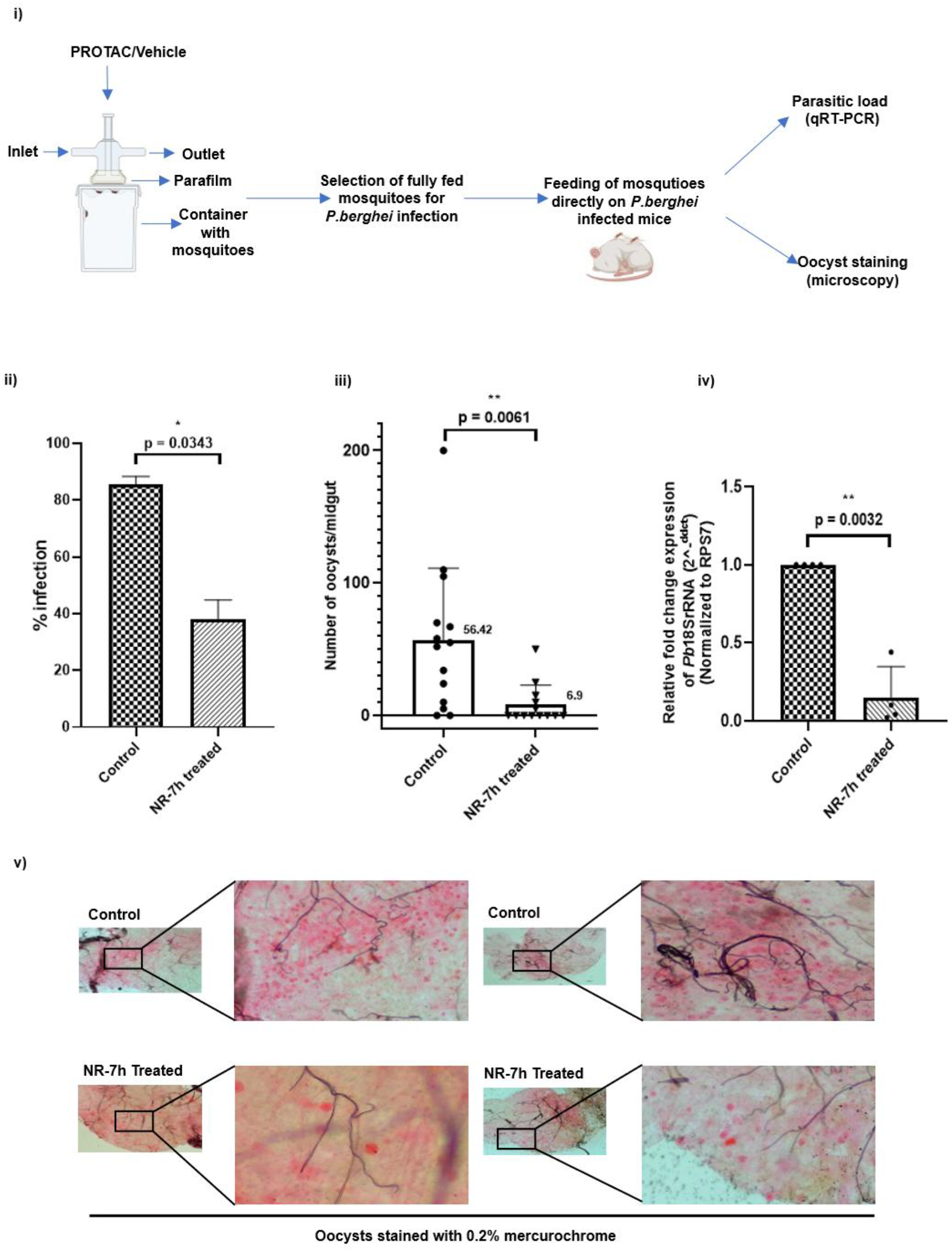
p38 MAPK degradation in *Anopheles* midgut inhibits *Plasmodium* development. (a) Schematic representation of the work flow was followed for determining the parasite growth and development under PROTAC treatment. PROTAC-mediated degradation of *As*p38 MAPK reduces (b) parasite infection, (c) mean oocyst count, and (d) parasitic load in midgut. (e) Microscopic images of 0.2% mercurochrome-stained midguts of control and treated mosquitoes.

### PROTACs modulates innate immunity to reduce parasite load in *Anopheles* midgut

A significant reduction in *Plasmodium* parasite load was observed following PROTAC treatment. To investigate the underlying mechanism, the expression levels of key innate immune response genes were analysed in the *Anopheles* midgut by qRT-PCR post 24 hours of *P. berghei* infection (24 h.p.i) under the effect of NR-7h treatment. Midgut expression of effector molecules including Defensin (ASTE011281), Gambicin (ASTEI0_032045), Cecropin C (ASTEI20_040486) and Nitric Oxide Synthase (NOS; ASTE008593) was analysed in dissected *P. berghei*-infected midguts following NR-7h treatment. Significant increase was observed in the expression of Defensin (∼35 fold), Gambicin (∼20 fold) and Cecropin C (∼50 fold), while a moderate increase was observed in the NOS (∼2 fold) expression (**Figure 5ai**). Additionally, the expression of immune signalling and regulatory components including Rel2 (ASTEI20_039998), TEP1 (ASTE016445), LRIM1 (ASTE000814), APL1 (ASTEI02571) and LRRD7/LRIM17 (ASTE009590) transcripts was analysed in dissected *P. berghei*-infected midguts following NR-7h treatment. Significant upregulation was observed in the transcript levels of LRRD7/LRIM17(∼5 fold), TEP1 (∼ 3-fold), LRIM1 (∼8 fold), APL1 (∼3 fold) and Rel2 (∼4 fold) in NR-7h treated *P. berghei*-infected group compared to the *P. berghei*-infected control group (**Figure 5a.ii**). These findings suggest that NR-7h treatment affects the mosquito’s ability to mount a defence during the initial parasite invasion of midgut epithelium by immediately triggering the innate immune responses. Similarly, the expression of the innate immune response genes in *Anopheles* midgut was also analysed post 8 days of *P. berghei* infection (8 d.p.i) to determine the effect of NR-7h mediated immune modulation on oocyst maturation thereby impacting the transmission potential. The results revealed that the expression profiles of the immune genes following NR-7h treatment remained comparable to that of 24 hours post-infection for both effector molecules **(Figure 5bi**) and immune signalling and regulatory components (**Figure 5bii**). The similar expression profile at both early and late stages of *P. berghei* infection suggest that NR-7h PROTAC exerts consistent and prolonged regulatory effect on mosquito midgut immune response. Collectively, these results indicate that NR-7h mediated *As*p38 MAPK degradation modulates the innate immune response in *Anopheles* during *Plasmodium* infection by the induction of immediate antimicrobial defence and activation of the complement-like and IMD pathways.

**Figure 5:**
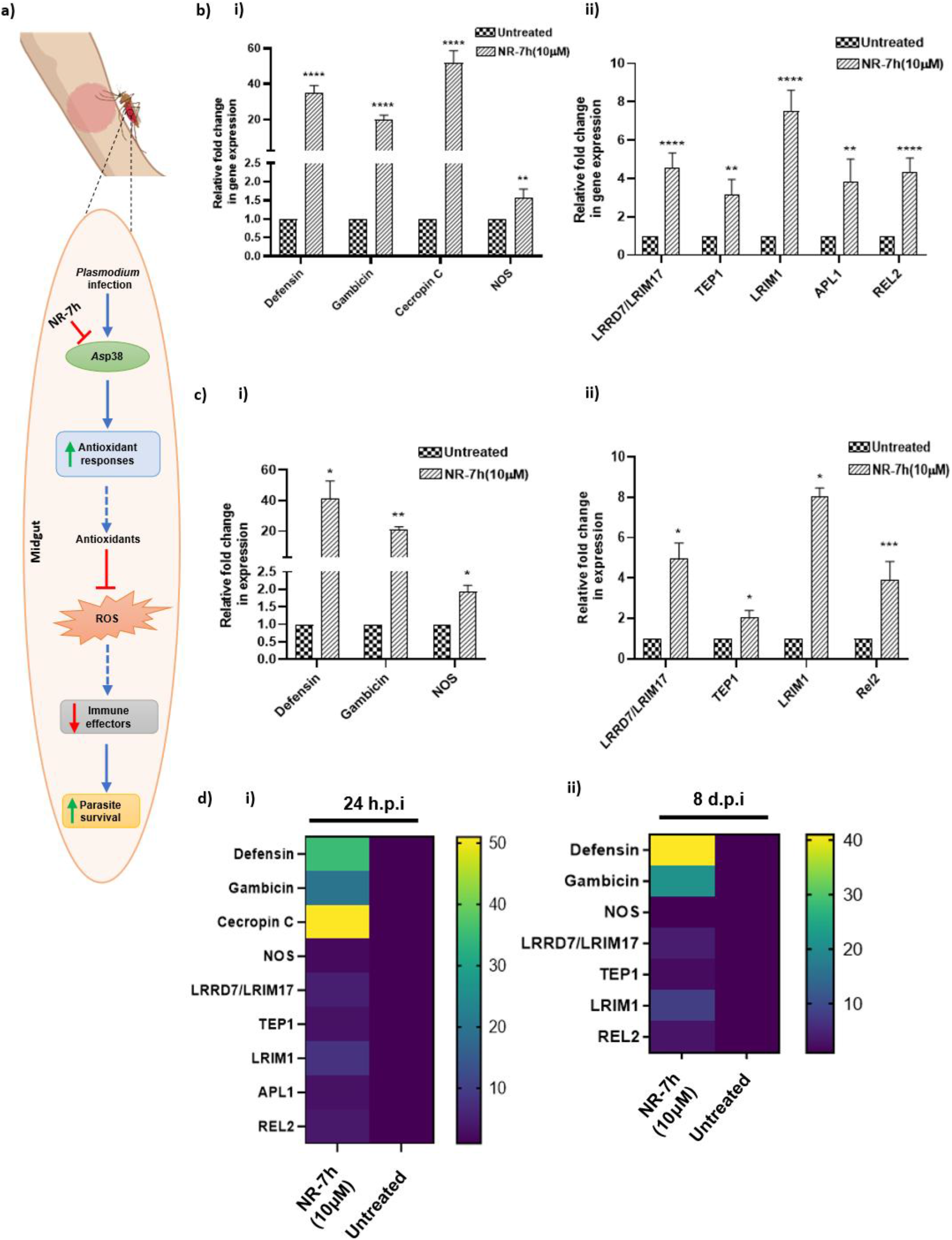
Effect of *As*p38 MAPK degradation on innate immune response. (a) Proposed model of NR-7h mediated negative regulation of *As*p38-dependent antioxidant regulation promoting *Plasmodium* survival in the *Anopheles* midgut. Expression of innate immune response genes in control and NR-7h treated dissected *Anopheles* midgut at (b) 24 hours-post infection of (i) antimicrobial defence genes and (ii) immune signalling and regulatory components and at (c) 8 days - post infection expression of (i) antimicrobial defence genes and (ii) immune signalling and regulatory components in control and NR-7h treated dissected *Anopheles* midgut. (d) Heat map representing the expression of innate immune response gene at (i) 24 hours post infection, and (ii) 8 days post infection.

## Discussion

Our study offers significant evidence that *Anopheles stephensi* has a functional Cullin-RING Ligase 4 (CRL4) complex and a conserved p38 MAPK signaling module, highlighting the evolutionary conservation of post-translational and signaling pathways in mosquitoes. The essential molecular systems governing protein turnover and stress signaling in vertebrates are also found to be conserved in vector mosquitoes, offering novel opportunities to investigate these pathways for targeting vector interventions.

Unlike the E3 ligases of humans, *Anopheles’* E3 ligases have not been extensively studied and characterized. For our study, the components of CRL4-CRBN complex, a Cullin Ring Ligand 4-E3 ubiquitin ligase complex, have been analysed in *A. stephensi*. The four main components of the complex include Cullin 4 (CUL-4), Ring-Box 1 (RBX-1), DNA Damage Binding protein1 (DDB1) and Cereblon (CRBN), where cereblon serve as the substrate receptor and is solely responsible for the binding to thalidomide and its derivatives used as E3 ligase binders in PROTACs^[52–54]^. Using the protein sequences of Human E3 ligase complex components, we could identify the components of CRL4-CRBN complex in *Anopheles stephensi*, which share significant similarity with Human homologues. *As*DDB1 (ASTEI20_032418), *As*RBX1(ASTE002617) and *As*CULLIN4B (ASTEI01623) share 67%, 84% and 64% similarity with human analogues, respectively. Human and *Anopheles* cereblon (*As*Cereblon; ASTEI20_034852) share relatively low similarity i.e. ∼37.2% identity at protein level, however the thalidomide binding domain has ∼46% identity along with the conservation of residues involved in binding to thalidomide and its analogues ^[54]^.

Evolutionary conservation of the CRL4–CRBN complex in *A. stephensi* has translational significance. In mammalian systems, this E3 ligase is reported to be hijacked by PROTACs to trigger selective target protein degradation^[49,50]^. Conserved residues in mammalian CRBN and *A. stephensi* imply that small molecules engineered to recruit human CRBN could also target the mosquito orthologue. This raises the intriguing possibility that PROTAC-like molecules could be optimized for protein degradation targeting in mosquito cells, creating a new platform for functional analysis and, ultimately, vector control through post-translational manipulation^[51,55]^.

Concurrently, our structural and phylogenetic characterization of *A. stephensi* p38 MAPK established high conservation of all key catalytic motifs—VAIK, HRD, DFG, and TXY—that are essential for activation of kinases and phosphorylation of substrates. From multiple sequence alignments to compare *Anopheles* homologues, we concluded that the substrate-binding and activation loop domains are closely related. This implies that the crucial regulatory and catalytic function has been conserved during evolution^[56,57]^. Phylogenetic reconstruction of *A. stephensi* p38 MAPK positioned it within the *Anopheles* clade, separate from Drosophila and vertebrate MAPKs, reflecting lineage-specific divergence while maintaining the kinase fold^[58]^. Structural superimposition on human p38 MAPK (RMSD = 0.642) confirmed the extensive three-dimensional conservation like the ATP-binding cleft and catalytic groove^[59]^.

Functional conservation of p38 MAPK among *Anopheles* species is most probably indicative of its pivotal function in stress and immune signaling. It modulates immune gene expression, oxidative stress regulation, and midgut epithelial integrity. In addition to modulate the generation of reactive oxygen species (ROS), which is critical for tissue homeostasis and parasite removal, p38 MAPK modulates the expression of several immunological effector molecules, such as detoxifying enzymes and antimicrobial peptides^[60]^. Research has demonstrated that disruption of *As*p38 MAPK signaling might hinder the development of oocysts, underscoring its possible function in promoting parasite survival and transmission competence.

Immune responses are major players in vector-pathogen interactions. Multiple immunological genes and pathways in *Anopheles stephensi* are essential for mediating anti-plasmodium responses and preserving vector competence^[61,62,63]^. The central to this defense is two pathways: Toll and Imd pathway to trigger the activation of two NF-κB transcription factors, Rel1 and Rel2^[64]^. This activation further led to the development of many effector molecules such as thioester-containing protein 1 (TEP1), fibrinogen-related proteins (FBNs), and antimicrobial peptides (AMPs)^[17,64,65]^. TEP1, which is stabilized by Leucine-rich Immune Molecule 1 (LRIM1) and Leucine-rich repeat protein 1 (APL1), is vital for identifying and removing *Plasmodium* ookinetes by lysis or melanization^[14,66]^. Additionally, NOS produces nitric oxide (NO), a reactive molecule with lethal effects on invasive parasites, which is regulated by immunological pathways such as Rel-2 and JAK/STAT^[20]^. Significantly, FREP1 (fibrinogen-related protein 1) and a conserved *Anopheles* mosquito prefoldin (PFDN)-chaperonin system were recently found to contribute to immune recognition, regulate protein homeostasis, and maintain midgut integrity during parasite invasion, all of which have an impact on infection outcomes^[63]^.

In this study, we also observed the activation of *As*p38 MAPK in the mosquito midgut upon infection with *Plasmodium* parasites. Since human p38 MAPK and *As*p38 MAPK share 72% of homology at protein level, we used a human-specific p38 PROTAC, NR-7h^[67]^ for the targeted degradation of *As*p38 MAPK, taking advantage of the evolutionary conservation of the p38 MAPK pathway across species. In contrast to traditional small-molecule inhibitors, which temporarily block kinase activity, PROTACs act by enabling the target protein to be selectively ubiquitinated, which leads to its breakdown through the proteasome pathway. This approach reduces the possibility of resistance development, which is frequently linked to active-site mutations, by degrading the kinase rather than simply inhibiting it. Here, we observed that PROTAC-mediated degradation of *As*p38 MAPK significantly reduces the parasite load in *Anopheles stephensi* mosquitoes’ midgut, confirming the crucial role of *As*p38 MAPK in malaria pathogenesis. According to studies, p38 MAPK helps maintain midgut homeostasis and epithelial repair resulting from parasite invasion by regulating the expression of detoxifying enzymes and antimicrobial peptides.

The regulation of immune-related genes and systems reflects a finely tuned interplay that determines vector competence. Therefore, the expression of Defensin, Gambicin, NOS, Rel2, LRRD7/LRIM17, TEP1, LRIM1 at the transcript level in NR-7h-treated and *P. berghei*-infected *Anopheles* midgut was also analysed. The increased expression of Defensin, Gambicin and NOS implies immediate induction of antimicrobial defence pathway while increase modulation of Rel2, LRRD7/LRIM17, TEP1, and LRIM1 reflects the longer-term immune readiness through pathogen recognition and signalling. Altogether, the modulation of innate immune response indicates the p38 MAPK-mediated regulation of these immune genes to favor parasite survival.

In summary, malaria transmission is entirely dependent on the mosquito vector, making this stage an attractive and critical target for therapeutic intervention. Our study reveals the conservation of the CRL4– CRBN E3 ligase complex and p38 MAPK signaling cascade in *A. stephensi*, illustrating a shared evolutionary framework of post-translational and stress-response mechanisms across taxa. By employing a human-derived PROTAC molecule to target *As*p38 MAPK, we demonstrate a novel strategy to disrupt parasite development and transmission. These findings open a translational avenue for exploiting conserved molecular machineries to design vector-stage interventions that complement existing anti-malarial strategies.

## Abbreviation

PROTAC: (PROtein Targeting Chimera)
MAPK: (Mitogen-Activated Protein Kinase)
*As*: (*Anopheles stephensi*)
(*Hs*): *Homo sapiens*
TEP1: (Thioester containing protein 1)
APL1: (Anopheles Plasmodium-responsive Leucine-rich repeat 1)
NOS: (Nitric Oxide Synthase)
LRIM1: (Leucine -Rich Immune Molecule 1)
LRRD7/LRIM17: (Leucine-Rich Repeat Domain-containing protein 7/ Leucine-Rich Immune Molecule 1)
Rel2: (Relish 2)

## Author contributions

S.S. has conceptualized the idea, designed the methodology, executed, analysed and curated the experimental data. S.S., K.P., and S.C. reviewed and edited the manuscript. N.R., J.S., and S.S. wrote the original draft of the manuscript. N.R., J.S., B.G., and N.P. designed the methodology. N.R., J.S., B.G., C.T., N.P., and KG. investigated the data. N.R., and J.S. performed formal analysis of the data. N.R., J.S., B.G. and N.P. validated the data and assisted in the data interpretation. All authors contributed to the article and approved the submitted version.

## Acknowledgements

We thank Prof. Angel R. Nebreda and Prof. Antoni Riera from the Institute for Research in Biomedicine, Barcelona for providing us with NR-7h, a PROTAC for p38-MAPK. We also thank the ICMR-National Institute of Malaria Research for providing the insectary facilities. We are grateful to JNU for providing access to the anti-plagiarism software Turnitin. We also thank Central Instrumentation Facilities of the Special Centre for Molecular Medicine, JNU. N.R. is supported by DBT-JRF. J.S. is a recipient of the BioCARe Women Scientist fellowship from DBT. S.S. is a recipient of the National Bio Scientist Award. This work has been funded by Indian Council of Medical Research project (IIRPSG-2024-01-03940) sanctioned to S.S.

## References

1. Guttery, D. S., Zeeshan, M., Ferguson, D. J. P., Holder, A. A. & Tewari, R. Division and Transmission: Malaria Parasite Development in the Mosquito. Annu. Rev. Microbiol. 76, 113–134 (2022).

2. Savi, M. K. medical sciences An Overview of Malaria Transmission Mechanisms, Control, and Modeling. Med Sci 11, (2023).

3. Siqueira-Neto, J. L. et al. Antimalarial drug discovery: progress and approaches. Nat. Rev. Drug Discov. 22, 807–826 (2023).

4. Thota, S. & Yerra, R. Drug Discovery and Development of Antimalarial Agents: Recent Advances. Curr. Protein Pept. Sci. 17, 275–279 (2016).

5. Clain, J., Hamza, A. & Ariey, F. Antimalarial Drugs for Malaria Elimination. Methods Mol. Biol. 2013, 151–162 (2019).

6. Laurens, M. B. Novel malaria vaccines. Hum. Vaccin. Immunother. 17, 4549–4552 (2021).

7. Beeson, J. G. et al. The RTS,S malaria vaccine: Current impact and foundation for the future. Sci. Transl. Med. 14, eabo6646 (2022).

8. Bevivino, G. et al. Peptides with Antimicrobial Activity in the Saliva of the Malaria Vector Anopheles coluzzii. Int. J. Mol. Sci. 25, (2024).

9. Kumar, S. & Barillas-Mury, C. Ookinete-induced midgut peroxidases detonate the time bomb in anopheline mosquitoes. Insect Biochem. Mol. Biol. 35, 721–727 (2005).

10. Baton, L. A. & Ranford-Cartwright, L. C. How do malaria ookinetes cross the mosquito midgut wall? Trends Parasitol. 21, 22–28 (2005).

11. Whitten, M. M. A., Shiao, S. H. & Levashina, E. A. Mosquito midguts and malaria: cell biology, compartmentalization and immunology. Parasite Immunol. 28, 121–130 (2006).

12. Vlachou, D. et al. Real-time, in vivo analysis of malaria ookinete locomotion and mosquito midgut invasion. Cell. Microbiol. 6, 671–685 (2004).

13. Povelones, M., Upton, L. M., Sala, K. A. & Christophides, G. K. Structure-function analysis of the anopheles gambiae LRIM1/APL1C complex and its interaction with complement C3-like protein TEP1. PLoS Pathog. 7, (2011).

14. Fraiture, M. et al. Two Mosquito LRR Proteins Function as Complement Control Factors in the TEP1-Mediated Killing of Plasmodium. Cell Host Microbe 5, 273–284 (2009).

15. Eleftherianos, I. et al. Haemocyte-mediated immunity in insects: Cells, processes and associated components in the fight against pathogens and parasites. Immunology 164, 401–432 (2021).

16. Lavine, M. D. & Strand, M. R. Insect hemocytes and their role in immunity. Insect Biochem. Mol. Biol. 32, 1295–1309 (2002).

17. Clayton, A. M., Dong, Y. & Dimopoulos, G. The Anopheles innate immune system in the defense against malaria infection. J. Innate Immun. 6, 169–181 (2014).

18. Saab, S. A., Cardoso-Jaime, V., Kefi, M. & Dimopoulos, G. Advances in the dissection of Anopheles–Plasmodium interactions. PLoS Pathog. 21, 1–30 (2025).

19. Horton, A. A. et al. The mitogen-activated protein kinome from Anopheles gambiae: identification, phylogeny and functional characterization of the ERK, JNK and p38 MAP kinases. BMC Genomics 12, 574 (2011).

20. Gupta, L. et al. The STAT pathway mediates late-phase immunity against Plasmodium in the mosquito Anopheles gambiae. Cell Host Microbe 5, 498–507 (2009).

21. Bondeson, D. P. et al. Catalytic in vivo protein knockdown by small-molecule PROTACs. Nat. Chem. Biol. 11, 611–617 (2015).

22. Evans, A. G. & Wellems, T. E. Coevolutionary genetics of Plasmodium malaria parasites and their human hosts. Integr. Comp. Biol. 42, 401–407 (2002).

23. Cuadrado, A. & Nebreda, A. R. Mechanisms and functions of p38 MAPK signalling. Biochem. J. 429, 403–417 (2010).

24. Wang, B. et al. Anopheles stephensi p38 MAPK signaling regulates innate immunity and bioenergetics during Plasmodium falciparum infection. Parasit. Vectors 8, 424 (2015).

25. Wang, L. et al. The tick saliva peptide HIDfsin2 promotes the tick-borne virus SFTSV replication in vitro by enhancing p38 signal pathway. Arch. Toxicol. 97, 1783–1794 (2023).

26. Bodhale, N. et al. Leishmania donovani mitogen-activated protein kinases as a host-parasite interaction interface. Cytokine 179, 156627 (2024).

27. Olson, C. M. et al. p38 mitogen-activated protein kinase controls NF-kappaB transcriptional activation and tumor necrosis factor alpha production through RelA phosphorylation mediated by mitogen- and stress-activated protein kinase 1 in response to Borrelia burgdorferi anti. Infect. Immun. 75, 270–277 (2007).

28. Kitsou, C. & Pal, U. Ixodes Immune Responses Against Lyme Disease Pathogens. Front. Cell. Infect. Microbiol. 8, 176 (2018).

29. Rolandelli, A. et al. Tick hemocytes have pleiotropic roles in microbial infection and arthropod fitness. bioRxiv Prepr. Serv. Biol. (2023) doi:10.1101/2023.08.31.555785.

30. Sakamoto, K. M. et al. Protacs: chimeric molecules that target proteins to the Skp1-Cullin-F box complex for ubiquitination and degradation. Proc. Natl. Acad. Sci. U. S. A. 98, 8554–8559 (2001).

31. Pettersson, M. & Crews, C. M. PROteolysis TArgeting Chimeras (PROTACs) - Past, present and future. Drug Discov. Today. Technol. 31, 15–27 (2019).

32. Churcher, I. Protac-Induced Protein Degradation in Drug Discovery: Breaking the Rules or Just Making New Ones? J. Med. Chem. 61, 444–452 (2018).

33. Burslem, G. M. & Crews, C. M. Proteolysis-Targeting Chimeras as Therapeutics and Tools for Biological Discovery. Cell 181, 102–114 (2020).

34. Sievers, F. & Higgins, D. G. Clustal Omega, Accurate Alignment of Very Large Numbers of Sequences. In Multiple Sequence Alignment Methods (ed. Russell, D. J.) 105–116 (Humana Press, 2014). doi:10.1007/978-1-62703-646-7_6.

35. Kumar, S. et al. MEGA12: Molecular Evolutionary Genetic Analysis Version 12 for Adaptive and Green Computing. Mol. Biol. Evol. 41, msae263 (2024).

36. Liu, Y. et al. CB-Dock: a web server for cavity detection-guided protein–ligand blind docking. Acta Pharmacol. Sin. 41, 138–144 (2020).

37. Sarikas, A., Hartmann, T. & Pan, Z.-Q. The cullin protein family. Genome Biol. 12, 220 (2011).

38. Rizvi, Z. et al. Plasmodium falciparum contains functional SCF and CRL4 ubiquitin E3 ligases, and CRL4 is critical for cell division and membrane integrity. PLoS Pathog. 20, e1012045 (2024).

39. Spratt, D. E., Wu, K., Kovacev, J.Pan, Z.-Q. & Shaw, G. S. Selective recruitment of an E2∼ubiquitin complex by an E3 ubiquitin ligase. J. Biol. Chem. 287, 17374–17385 (2012).

40. Zheng, N. et al. Structure of the Cul1-Rbx1-Skp1-F boxSkp2 SCF ubiquitin ligase complex. Nature 416, 703–709 (2002).

41. Nguyen, H. C., Yang, H., Fribourgh, J. L., Wolfe, L. S. & Xiong, Y. Insights into Cullin-RING E3 ubiquitin ligase recruitment: structure of the VHL-EloBC-Cul2 complex. Structure 23, 441–449 (2015).

42. Lee, J. & Zhou, P. DCAFs, the missing link of the CUL4-DDB1 ubiquitin ligase. Mol. Cell 26, 775– 780 (2007).

43. Stahl, K. et al. Modelling protein complexes with crosslinking mass spectrometry and deep learning. Nat. Commun. 15, 7866 (2024).

44. Hammarén, H. M., Virtanen, A. T. & Silvennoinen, O. Nucleotide-binding mechanisms in pseudokinases. Biosci. Rep. 36, e00282 (2015).

45. La Sala, G. et al. HRD Motif as the Central Hub of the Signaling Network for Activation Loop Autophosphorylation in Abl Kinase. J. Chem. Theory Comput. 12, 5563–5574 (2016).

46. Filomia, F., De Rienzo, F. & Menziani, M. C. Insights into MAPK p38alpha DFG flip mechanism by accelerated molecular dynamics. Bioorg. Med. Chem. 18, 6805–6812 (2010).

47. Zarubin, T. & Han, J. Activation and signaling of the p38 MAP kinase pathway. Cell Res. 15, 11–18 (2005).

48. Han, J., Wu, J. & Silke, J. An overview of mammalian p38 mitogen-activated protein kinases, central regulators of cell stress and receptor signaling. F1000Research 9, (2020).

49. Ito, T., Yamaguchi, Y. & Handa, H. Exploiting ubiquitin ligase cereblon as a target for small-molecule compounds in medicine and chemical biology. Cell Chem. Biol. 28, 987–999 (2021).

50. Baek, K. et al. Unveiling the hidden interactome of CRBN molecular glues. Nat. Commun. 16, 6831 (2025).

51. Nguyen, T. Van. USP15 antagonizes CRL4(CRBN)-mediated ubiquitylation of glutamine synthetase and neosubstrates. Proc. Natl. Acad. Sci. U. S. A. 118, (2021).

52. Fischer, E. S. et al. Structure of the DDB1-CRBN E3 ubiquitin ligase in complex with thalidomide. Nature 512, 49–53 (2014).

53. Mori, T. et al. Structural basis of thalidomide enantiomer binding to cereblon. Sci. Rep. 8, 1294 (2018).

54. Yamamoto, J., Ito, T., Yamaguchi, Y. & Handa, H. Discovery of CRBN as a target of thalidomide: a breakthrough for progress in the development of protein degraders. Chem. Soc. Rev. 51, 6234– 6250 (2022).

55. Rosignoli, S., Giordani, S., Pacelli, M., Guarguaglini, G. & Paiardini, A. Unraveling the Engagement of Kinases to CRBN Through a Shared Structural Motif to Optimize PROTACs Efficacy. Biomolecules 15, (2025).

56. Guo, Z. et al. The regulation landscape of MAPK signaling cascade for thwarting Bacillus thuringiensis infection in an insect host. PLoS Pathog. 17, e1009917 (2021).

57. Yao, M., Liu, X., Meng, J., Yang, C. & Zhang, C. Molecular characterization and elucidation of the function of Hap38 MAPK in the response of Helicoverpa armigera (Hübner) to UV-A stress. Sci. Rep. 12, 18489 (2022).

58. Han, Z. S. et al. A conserved p38 mitogen-activated protein kinase pathway regulates Drosophila immunity gene expression. Mol. Cell. Biol. 18, 3527–3539 (1998).

59. Wilson, K. P. et al. Crystal structure of p38 mitogen-activated protein kinase. J. Biol. Chem. 271, 27696–27700 (1996).

60. Surachetpong, W., Pakpour, N., Cheung, K. W. & Luckhart, S. Reactive oxygen species-dependent cell signaling regulates the mosquito immune response to Plasmodium falciparum. Antioxid. Redox Signal. 14, 943–955 (2011).

61. Kajla, M. et al. Anopheles stephensi Heme Peroxidase HPX15 Suppresses Midgut Immunity to Support Plasmodium Development. Front. Immunol. 8, 249 (2017).

62. Sharma, P. et al. Altered Gut Microbiota and Immunity Defines Plasmodium vivax Survival in Anopheles stephensi. Front. Immunol. 11, 609 (2020).

63. Dong, Y. et al. Targeting the mosquito prefoldin-chaperonin complex blocks Plasmodium transmission. Nat. Microbiol. 10, 841–854 (2025).

64. Zakovic, S. & Levashina, E. A. NF-κB-Like Signaling Pathway REL2 in Immune Defenses of the Malaria Vector Anopheles gambiae. Front. Cell. Infect. Microbiol. 7, 258 (2017).

65. Garver, L. S. et al. Anopheles Imd pathway factors and effectors in infection intensity-dependent anti-Plasmodium action. PLoS Pathog. 8, e1002737 (2012).

66. Waterhouse, R. M., Povelones, M. & Christophides, G. K. Sequence-structure-function relations of the mosquito leucine-rich repeat immune proteins. BMC Genomics 11, 531 (2010).

67. Singhal, J. Chemical degradation identified Human p38-MAPK as a host target to combat parasitic infections. (2024).

